# Classification of Free-Living Body Posture with ECG Patch Accelerometers: Application to the Multicenter AIDS Cohort Study

**DOI:** 10.1101/2021.07.16.452654

**Authors:** Lacey H. Etzkorn, Amir S. Heravi, Katherine C. Wu, Wendy S. Post, Jacek Urbanek, Ciprian Crainiceanu

## Abstract

As health studies increasingly monitor free-living heart performance via ECG patches with accelerometers, researchers will seek to investigate cardio-electrical responses to physical activity and sedentary behavior, increasing demand for fast, scalable methods to process accelerometer data. We provide the first published analysis of tri-axial accelerometry data from Zio XT patch and introduce an extension of posture classification algorithms for use with ECG patches worn in the free-living environment. Our novel extensions to posture classification include (1) estimation of an upright posture for each individual without the reference measurements used by existing posture classification algorithms; (2) correction for device removal and re-positioning using novel spherical change-point detection; and (3) classification of upright and recumbent periods using a clustering and voting process rather than a simple inclination threshold used in other algorithms. Methods were built using data from 14 participants from the Multicenter AIDS Cohort Study (MACS), and applied to 1, 250 MACS participants. As no posture labels exist in the free-living environment, we evaluate the algorithm against labelled data from the Towson Accelerometer Study and against data labelled by hand from the MACS study.

## 1. Introduction

The Zio XT patch (Zio) is an ambulatory waterproof single-channel electrocardiographic (ECG) monitor that is FDA approved to record a person’s cardiac electrical rhythm for up to two weeks during daily life (Zio, iRhythm Technologies Inc., San Francisco, Calif). While Zio is primarily marketed as tool to diagnose rare cardiac arrythmias, Zio has been used by researchers to record ECG features over a two week period, quantify these features’ circadian and ultradian patterns, and analyze their association with health conditions and outcomes.

Zio was used in a substudy of the Multicenter AIDS Cohort Study (MACS) to explore whether men living with HIV experience abnormal ventricular repolarization variability, a known correlate of sudden cardiac death in HIV uninfected populations (Deyell *and others*, 2015). The MACS substudy used Zio ECG data to derive participants’ QT Variability Index, a measure of abnormal ventricular repolarization, and examine associations with HIV (Heravi *and others*, tted). However, MACS researchers needed to adjust for differences in time spent recumbent by HIV serostatus, as QT Variability Index is known to change with body posture(Yeragani *and others*, 2000a).

As Zio also contains a tri-axial accelerometer, we extracted and processed the accelerometry data from the files provided by Zio and developed an approach for classifying whether a person is recumbent or upright at a given time. Our work identifies and addresses some challenges for classifying body posture from ECG patch accelerometers worn in the free-living environment that previous work on posture classification has not addressed. These challenges include (1) estimating reference postures without time-intensive lab measurements that other algorithms rely on; (2) accounting for intermittent device repositioning; and (3) addressing ‘‘speckling” in posture classification when an individual’s posture is near the boundary between upright and recumbent. Additionally, to the best of our knowledge, this is the first time that Zio accelerometer measurements have been processed and used in a study. Having the accelerometer data in addition to the ECG data from Zio provides opportunities for researchers to study cardiac response to physical activity and posture.

In section 2.2 we demonstrate how posture can be visibly recognized in Zio accelerometer data, and we build the intuition behind our approach for posture classification. In Section 3 we (1) describe existing approaches for posture classification from accelerometer data, (2) introduce additional challenges for posture classification with ECG monitors in large cohort studies, and (3) detail our solutions to these unique challenges using both equations and visualizations. Section 4 provides the results of the algorithm applied to Zio data from the MACS substudy and to Actigraph data from the Towson Accelerometer Study. Section 5 provides a discussion of results and their potential limitations. The appendix includes information about our software for posture classification, a software vignette, and sensitivity analyses for our algorithm’s four tuning parameters.

## 2. Data

### 2.1 MACS Study and Sample

Zio ECG and tri-axial accelerometer data were obtained for 1, 250 men from MACS. MACS is a cohort study of men living with and without HIV infection in Baltimore, Chicago, Pittsburgh, and Los Angeles. After a biannual visit in 2016 and 2017, MACS participants could elect to wear Zio for up to 14 days. Methods for recumbent estimation were developed using data on 14 MACS participants with Zio accelerometry data and the resulting algorithm was applied to the 1, 250 men from the MACS study.

### 2.2 Accelerometer Data: Background and Features

The top panel in Figure 1 provides an example of Zio accelerometer data over the course of one day (midnight to midnight) for one person. The bottom panel in the same figure displays concurrent heart rate data for the same study participant.

**Fig. 1.**
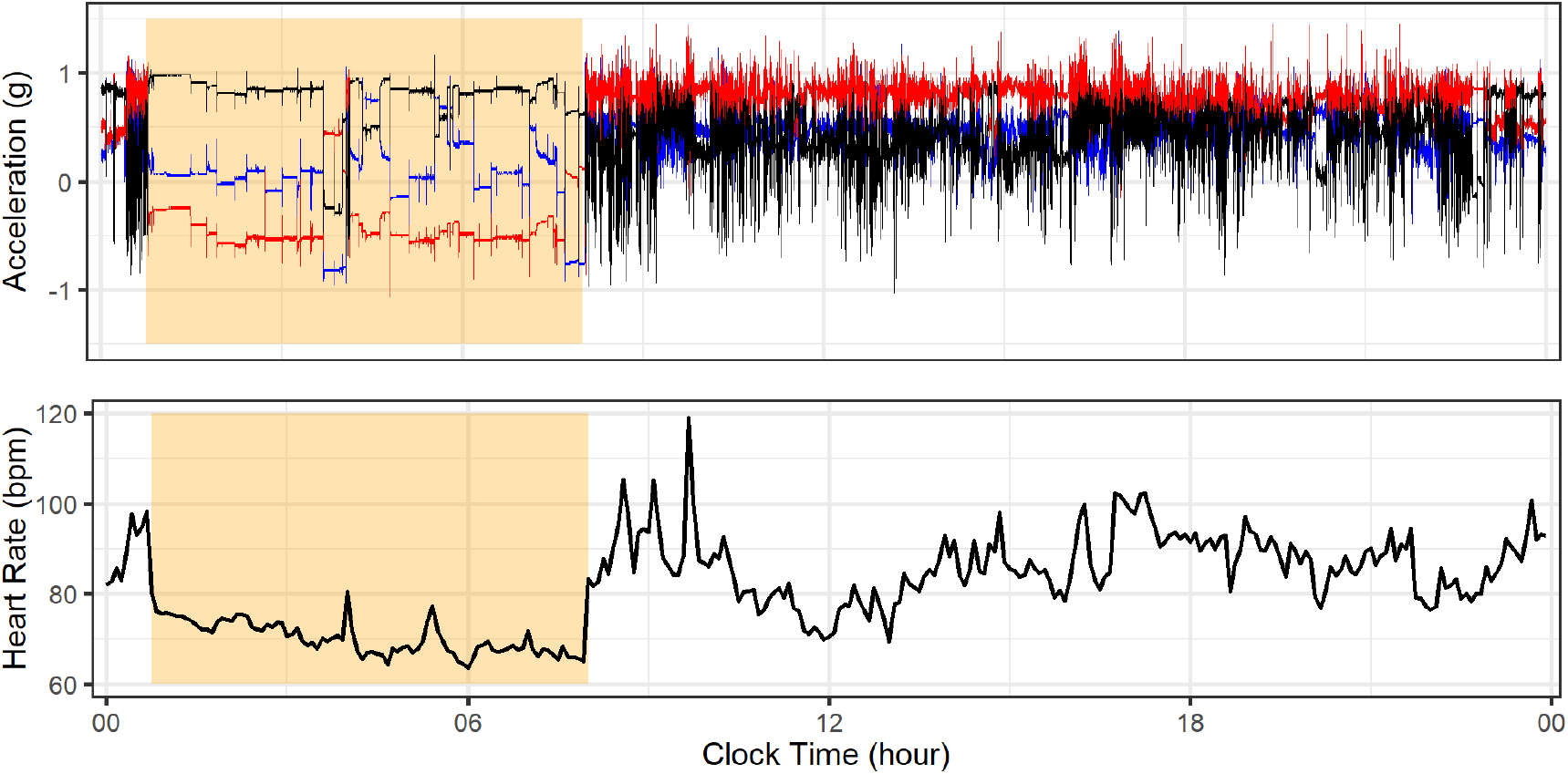
Top: 24 hours of Zio triaxial accelerometry for a MACS subject. Accelerations along the X (blue), Y (red), and Z (black) axes, beginning at midnight. Bottom: ECG-Derived Heart Rate in 5 minute intervals. The orange-highlighted regions correspond to an area of interest discussed in the text.

At the most basic level, accelerometers measure acceleration in gravitational units (*g*) along three orthogonal axes (back/forward, up/down, left/right) with respect to the accelerometer’s frame of reference. Measurements from these three axes are shown as black, red, and blue timeseries in Figure 1. Because Zio is affixed to the chest, the observed accelerometry is a reasonable proxy for the acceleration of the chest. The top panel in Figure 1 shows distinct differences in the tri-axial acceleromery signal between 1am to 8am (highlighted in orange) versus the period 8am to 10pm. The ordering of the three axes data is changed during the 1am to 8am period, with the axis shown in black hovering just below 1*g*, the axis shown in red hovering around —0.5*g*, and the axis shown in blue being less stable but having an overall mean close to 0*g*. Moreover, the amplitude of the signal is much lower for each axis, containing long periods with few changes in relative size of the signals. In contrast, the ordering of the axes changes during the 8am to 10pm period and the amplitude of the signal is substantially larger. Indeed, the axis shown in red is hovering close to 1*g*, while the axis shown in black is hovering slightly above 0.25*g*. Moreover, the distinct differences in the signals between the two periods are reflected in the heart rate, shown in the bottom panel. The heart rate is markedly lower during the 1am to 8am period.

These differences in the accelerometry patterns are consistent with a difference in the inclination of the chest. As the Earth’s gravitational force acts continuously on the accelerometer, it pulls the device with 1*g* acceleration towards the center of the earth. Thus, when the device changes position relative to Earth’s gravitation, the device’s axes change alignment with respect to gravity, which changes the relative acceleration measured along the three accelerometer axes. The change in relative acceleration corresponds to the reordering of the axes seen in the top panel of Figure 1. For example, in the top panel of Figure 1 at 6am, the device measures (0.09, —0.52, 0.85)*g* along the X (blue), Y (red), and Z (black) axes, respectively. At noon, the device measures (0.51, 0.80, 0.32)*g*. The angular difference between these two orientations of the device is

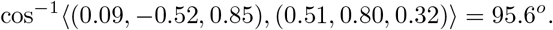

This indicates that at 6am the the chest is rotated 95 degrees relative to its position at 12pm. This suggests that the individual was upright at one of these time points and recumbent at the other.

But at which of these time points is the participant upright? We believe the answer lies in the relative intensity of physical activity performed during these time periods. Previous work has demonstrated that the amplitude of acceleration along each axis is highly correlated with activity intensity (Karas *and others*, 2019). In general, more intense physical activity will induce more accelerations and decelerations on the accelerometer. Consequently, the lower amplitude along each axis in the orange-highlighted region implies that the person is moving far less during this period. Hence the orange-highlighted period is more likely to represent a recumbent state. The fact that the heart rate decreases substantially in the 1am to 8am period is also consistent with a period of being recumbent.

In summary, recumbent rest exhibits lower amplitude accelerations along the three axes and a change in the relative ordering of the axes relative to periods with high amplitude of acceleration. In the next section we outline how to leverage these phenomenon to classify posture.

## 3. Challenges and Solutions for Posture Classification with Zio

Using triaxial accelerometers to classify posture is not new. For example, (Hansson *and others*, 2001) start with a reference upright orientation vector *μ_i_*; ||*μ_i_*|| = 1 for each person *i* = 1,…, *n*. This is a three-dimensional unit vector that represents the reading of the tri-axial accelerometer when a person stands upright and still. Given the measurement of the device *Y_i_*(*t*) at any time *t* = 1,…, *T_i_* the angle of inclination can be estimated as *θ_i_*(*t*) = cos^-1^(〈*μ_i_*, *Y_i_*(*t*)/||*Y_i_*(*t*)||〉). Last, a decision rule, such as *θ_i_*(*t*) > π/4, is used to label whether a person is recumbent or upright. Variations of this general approach are used by different platforms to estimate inclination and classify posture. For example, the ActiLife software classifies posture using ActiGraph accelerometry data when the device is mounted at the waist by assuming that *μ_i_* = (0, — 1,0) and estimating that a person is recumbent if the inclination angle is greater than 65 degrees (*θ_i_*(*t*) > 1.13) (ActiGraph, 2018).

Despite the existing general framework for classifying posture, implementing this framework for Zio accelerometer data in the MACS study raises several methodological problems. First, the upright posture, *μ_i_*, must be estimated for each person, as a standardized upright posture were not recorded for any of the study participants. Furthermore, *μ_i_* may need to be time-varying if participants remove and reposition Zio on their chests. Second, the decision rule for classifying upright and recumbent postures may need to be participant-specific, as both the inclination and range of activities may vary substantially across study participants. Third, algorithm performance must be evaluated in the absence of a gold-standard. We detail our approaches for these challenges in Sections 3.1 through 3.4

### 3.1 Estimate the upright posture

We start with the assumption that individuals tend to perform their most vigorous physical activities while upright. Therefore, one could expect that most intense activities (as measured by acceleration) are centered around the upright position vector. To illustrate the concept, the top row in Figure 2 displays the triaxial accelerometer measurements in 3D space for three study participants.

**Fig. 2.**
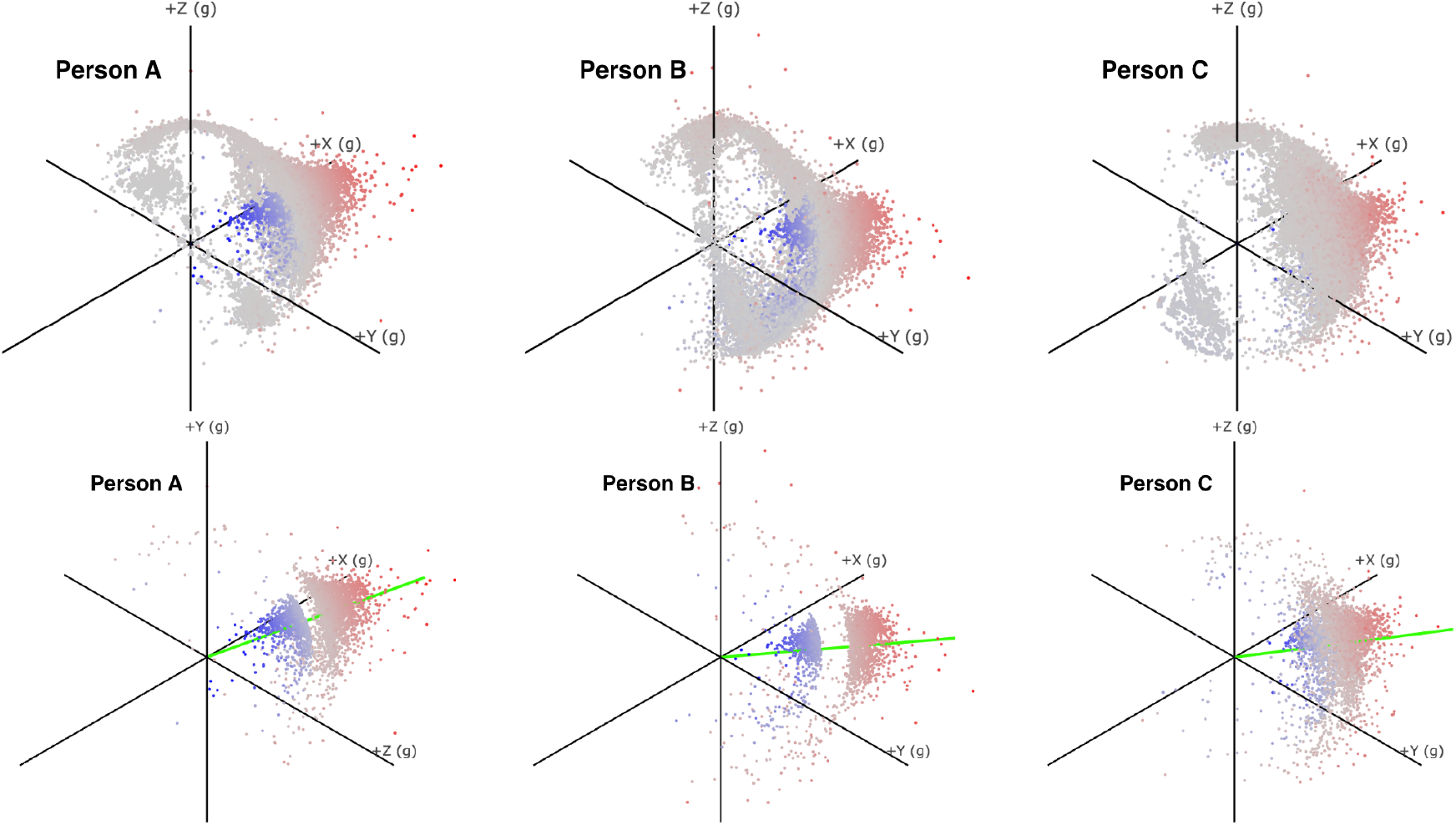
Top row: All Zio tri-axial accelerometry measurement in 3D space for three participants. Deeper shades of red correspond to vector magnitudes (*L*_2_ norm) that are larger and farther away from 1*g* (outside the unit sphere). Deeper shades of blue correspond to vector magnitudes (*L*_2_ norm) that are smaller and farther away from 1g (inside the unit sphere). Points shown in gray are close to the unit sphere. Bottom row: Same plots as above, but with the gray dots removed. Green lines represent the central orientation of high acceleration points.

Figure 2 indicates that, as expected, high acceleration periods tend to cluster in one location on the unit sphere for each individual; note the deeper shades of red and blue dots. At rest, the vector magnitude of the tri-axial accelerometer measures 1g (or close to 1g), and lies close to the unit sphere. The gray points in the top panel of Figure 2 are the points that correspond to no- or very low-intensity activity. In contrast, when individuals are active, the vector magnitude tends to be farther away from the surface of the sphere, as indicated by the red and blue dots; deeper shades of color correspond to larger distances from those points to the unit sphere. We focus on estimating the central vector orientation for each individual among the high acceleration vectors. For individual *i* at time *t*, denote the time series of tri-axial accelerations measured at 1.56Hz (approximately 94 samples/minute) as *X_i_*(*t*) = {*x_i_*(*t*), *y_i_*(*t*), *z_i_*(*t*)} and define *r_i_*(*t*) = ||*X_i_*(*t*)|| for *t* = 1,…, *T_i_* the vector magnitude of X_i_(t). For each study participant, the approach first identifies the times *H_i_* when acceleration is above their *p* = 0.95 empirical quantile:

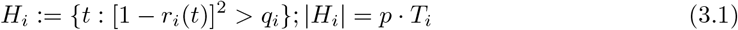

The default value of the parameter *p*\* we use to define *H_i_* was chosen using a sensitivity analysis further described in Appendices 2a and 2b. For each individual, {*X_i_*(*t*): *t* ∈ *H_i_*} is displayed in the bottom row of Figure 2. Once H_i_ is extracted, the mean direction 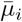 and mean resultant length *R_i_* of *H_i_* are defined as

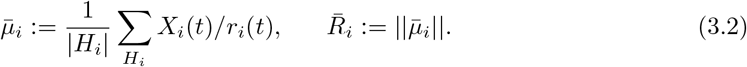

The estimated upright position 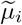 is the mean direction of *H_i_* scaled to unit length:

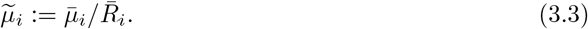

The estimated upright orientations are displayed as green vectors in Figure 2.

### 3.2 Detect device removal and re-position

The estimation procedure in the previous section depends on the assumption that the device was not removed and re-positioned. Re-positioning of the device could substantially alter the orientation of the device when a person is upright. Close inspection of the data indicated that for three of sixteen study participants in the training set (18.8%), two distinct clusters of high activity could correspond to the upright posture. Figure 3 displays the tri-axial measurements plotted in 3D space for two individuals. The data for Person 1 in the left panel displays two concentrated orientations of high-intensity activity (note the red dots clustering at the opposite poles of the sphere). This may indicate that the participant removed the device and re-positioned it upside-down. The data on the right for Person 2 indicates only one cluster of high intensity activity. This is likely consistent with no device removal.

**Fig. 3.**
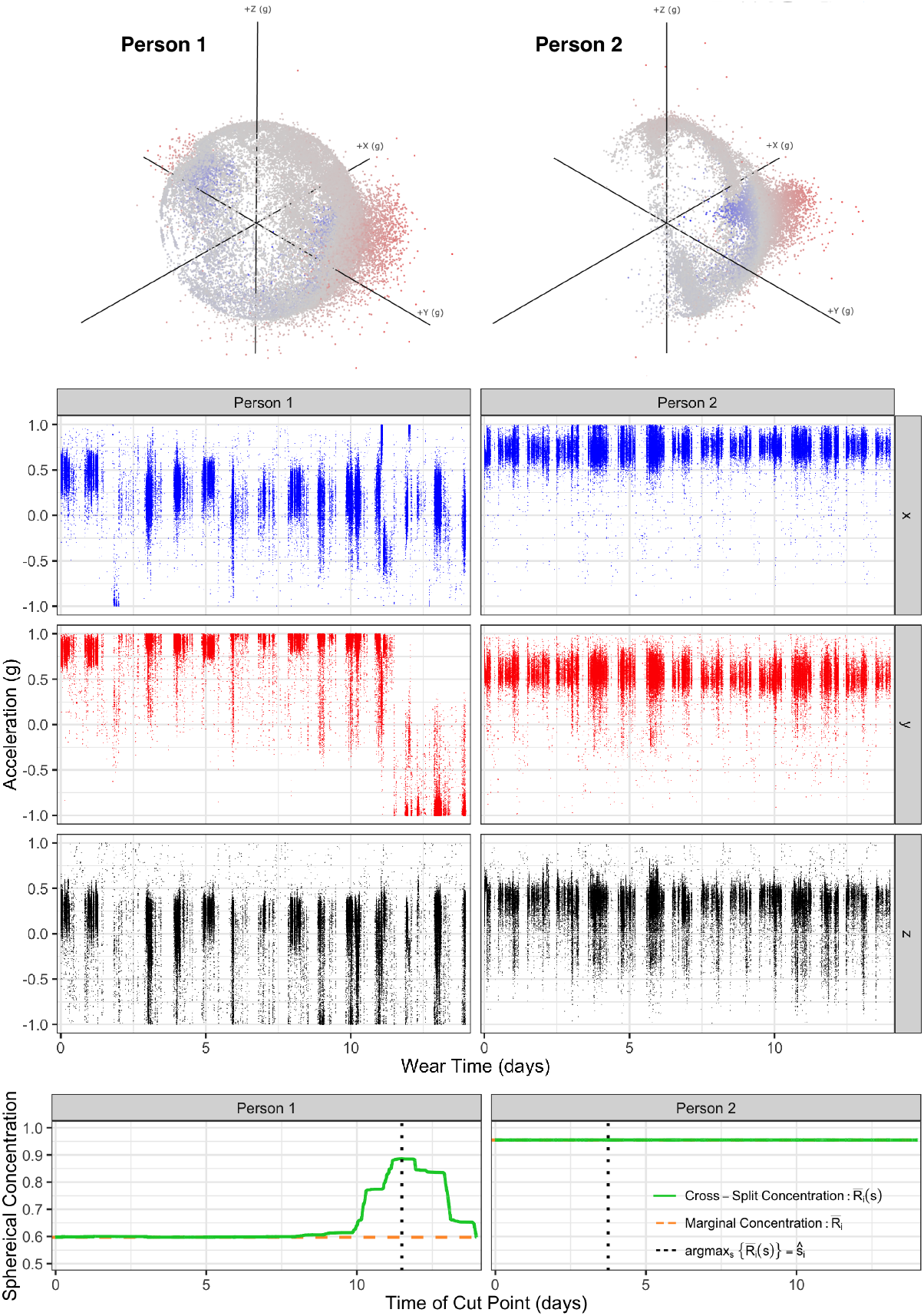
Top Panels: Highest accelerations in 3D space for two individuals. Center Panels: Acceleration time series for the same two individuals across the three axes. Bottom Panels: Marginal and cross-split concentration of high-acceleration points. The vertical dashed line indicates the point where the cross-split angular concentration is maximized.

The three panels below the sphere for Person 1 display the three acceleration time series shown only for the top 5% highest accelerations: {*X_i_*(*t*): *t* ∈ *H_i_*}. Closer inspection of the time series shown in red indicates that a clear change occurred around day 11 (note the mean shift from around 0.9g to —0.9g). An examination of the ECG data displayed a flip in the wave forms around day 11, which further re-enforces the idea that the device was removed and flipped nearly 180 degrees.

Therefore, we complement the approach in the previous section with a change point approach that: (1) detects whether an individual has removed and re-positioned the device; and (2) estimates the time when this happened. We examine a measure of combined spherical concentration before and after every potential change point to estimate whether and when the device was removed. If a change in orientation exists then one would expect to see a high spherical concentration within each split of the data relative to the overall marginal concentration.

We examine every time point *s* ∈ *H_i_* and denote *n_bi_* (*s*) and *n_ai_* (*s*) the number of high-acceleration observations before and after *s*; the total number of high-acceleration observations for study participant *i* is |*H_i_* | = *n_bi_*(*s*) + *n_ai_*(*s*) + 1. We estimate the mean direction, mean resultant length, and upright orientation before (*b*) and after (*a*) time *s* as

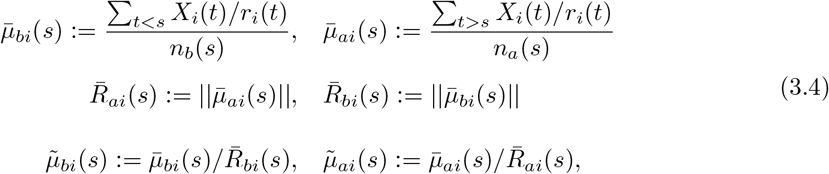

The cross-split mean resultant length is the weighted average of mean resultant lengths before and after time *s*:

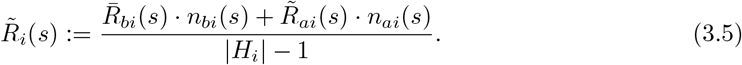

The time 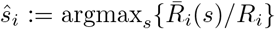 is when the max cross-split mean resultant length, 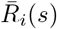, is maximized. The bottom panels of Figure 3 display the cross-split mean resultant length 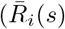, solid) as a function of time. The marginal mean resultant length, 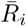, is represented as a horizontal, dashed line and the time of the maximal cross-split mean resultant length (*ŝ_i_*) is the vertical, dotted line.

Figure 3 demonstrates that these measures capture the timing and presence of the change that was visually identified to be around day 11 for person 1 in the central panel. For person 1, the cross-split mean resultant length is flat for the first 10 days, it starts increasing and attains its maximum sometimes during day 11, and then decreases. In contrast, for person 2 all mean resultant length are nearly equal, indicating there is little evidence the upright position for person 2 changed at any point in time.

Figure 3 also demonstrates that the marginal mean resultant length for person 1 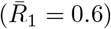 is much lower than for person 2 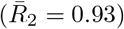. This makes sense because the high acceleration points for person 1 are dispersed across two clusters, whereas for person 2 they are mean resultant length in one cluster. However, cutting the data at time 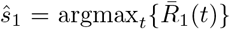 increases the cross-split mean resultant length for person 1 to a much larger mean resultant length, 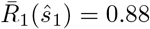.

We propose to use the ratio 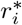 of the maximal cross-split mean resultant length and marginal mean resultant length 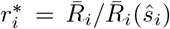 to determine whether the device was removed and repositioned. The use of the cross-split mean resultant length in a change point analysis was inspired by the use of a similar statistic in two-sample tests for differences in central orientations of spherical data (Mardia and Jupp, 2000). If there is a true difference in central orientation between two splits, then the cross-split mean resultant length will be high relative to the marginal mean resultant length, and 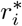 will be closer to 0. As in all changepoint analyses, the two sample test cannot be used directly because of (1) autocorrelation in the samples and (2) inference is performed after the cross-split concentration has been maximized. For a formal test, a cutoff *k* for the ratio 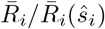 should be chosen to ensure that the associated statistical test has a particular size, say *α* = 0.05. However, for our purposes, we choose *k* = 0.98, as this seemed to demarcate participants whose data displayed visual evidence of device removal (Appendix 2a). The three participants in the test set of 16 whose devices seemed to be removed had ratios of 0.67 to 0.91, while the 13 participants who did not display visual evidence of device removal had ratios ranging from 0.9908 to 0.9999. Therefore, we evaluate the sensitivity of our results to various cutoffs in Appendix 2a.

### 3.3 Classify Intervals of Recumbent Rest

After estimating the upright orientation for each individual, 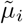, we proceed with estimating body orientation in 5-minute intervals, though we found our algorithm to work well for both 1 minute intervals and 30 second intervals as well. Let *t_j_*; *j* = 1,…,*J_i_* be the sequence of times indexing the 5-minute time intervals. For two weeks of recording time, *J_i_* = 3, 456. Our estimate of the orientation of the device *Ŷ_i_*(*t*), inclination of the chest 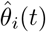, and initial indicator for upright posture 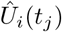 at time *t* in the *j*th interval are

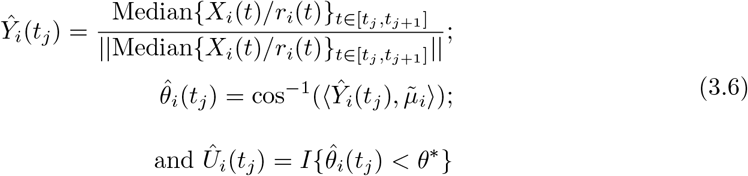

where 〈.,.〉 denotes the inner product and ||.|| denotes the *L*_2_ norm. We set *θ*\* = π/4 as a default value, as it marks the half way point between a perfectly upright position (0) and perfectly recumbent postition (π/2), though we examine the sensitivity of classifications to *θ*\* in Appendix 2d. The results of this initial classification are illustrated in the top two rows of Figure 4 for three study participants, labeled A, B, and C, the same study participants shown in Figure 2 from Section 3.1. The first and second row of Figure 4 provide two 2D perspectives of the participant-specific sphere (each participant is shown in one column) together with the estimated median orientations, *Ŷ_i_*(*t*), in 5-minute intervals. In the top row the circles mark 45 and 90 degrees of inclination from the estimated upright orientation and the “×” at the center marks 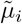. The second row shows the same data from a different perspective, with the lower and upper lines corresponding to the same 45 and 90 degrees of inclination. Red color indicates an initial label of upright posture: 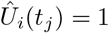.

**Fig. 4.**
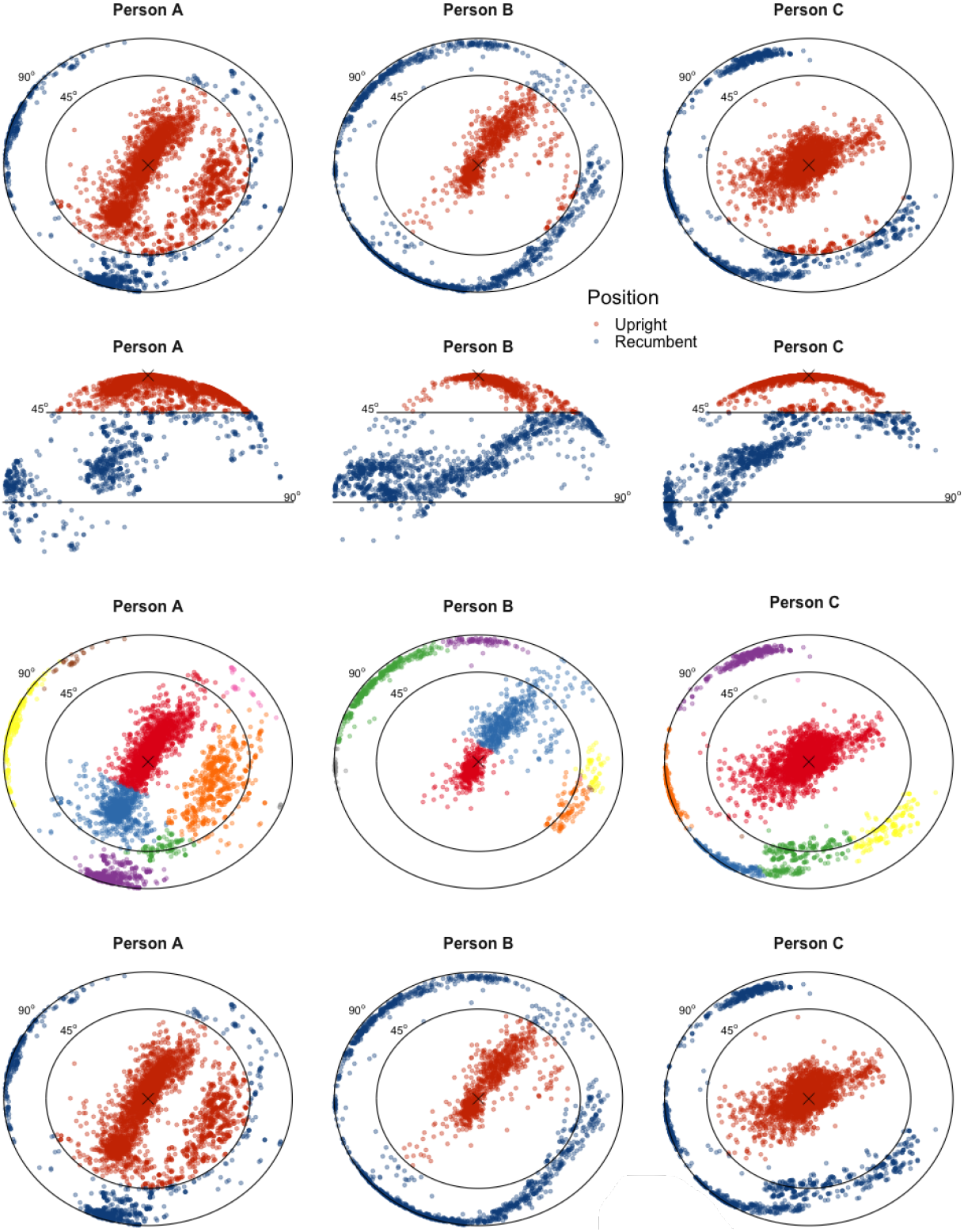
Each point represents the median orientation of the device for each five-minute interval for three individuals. The symbol × in each plot corresponds to the individual’s upright orientation. Each person’s data has been rotated toward a common upright posture. Row 1 and 2: Angular-threshold labels for recumbent. Row 3: Results of mean-shift clustering (bandwidth = 0.14). Row 4: Cluster-refined labels for recumbent posture.

Each individual’s upright position is centered in an oblong cluster, which likely represents the variation within each person’s upright position. The cluster is probably oblong because the backward/forward deviations from the upright position are much more common than left/right deviations while being upright. Clusters forming girdles around the sides of the spheres likely represent classes of recumbent positions for each individual. A closer inspection of the top two panels in Figure 4 indicates that there might be some problems with the initial classification approach of the median inclinations, *Y_i_*(*t*). For example, for each of the three study participants there are clusters that have some points classified as upright and some points classified as recumbent. The problem seems to occur at any level of angular thresholding, so simply changing the threshold from 45 to 40 degrees may not present a more acceptable solution. We propose to use this clustering information to refine the classification algorithm. More precisely, we use a clustering saturation approach, where we cluster the data in a large number of clusters and then, we estimate all points in the cluster as being either upright or recumbent using a majority rule voting. That is, if more than 50% of the points in the cluster were estimated in the initial classification to be recumbent then all points in the cluster are estimated to be recumbent.

We visually investigated the results of several clustering approaches for the 14 study participants in our exploratory sample. The choice of the clustering algorithm and sensitivity of results to this choice are detailed in Appendices 1 and 2d, respectively. All clustering approaches assigned the median orientations, *Ŷ_i_*(*t_j_*), into *K_i_* clusters *G_ik_*, *k* = 1… *K_i_*. The results of the clustering approach on the three study participants in Figure 4 are shown in the bottom two rows of the same figure. In the third row each individual color indicates an estimated cluster (i.e. *j* ∈ *G_ik_*).

We refine the initial indicators for an upright position 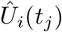 by re-labelling each point recumbent if the majority of time intervals in it’s cluster were labelled recumbent:

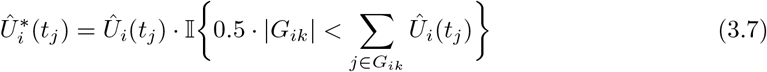

The fourth row in Figure 4 provides the final labels, which can be compared to the initial estimates in the second panel. One can see that around the decision area, π/4, the clusters are now in one color, which is more reasonable given the proximity of the points.

### 3.4 Quality Control

We cannot validate our approach using internal gold-standard labels of upright and recumbent states with data from the MACS study since participants wore Zio in the free-living environment. Instead, we validate our algorithm with manual inspection of the classifications as well as validation against labelled data collected in a lab in a slightly different setting.

First we examine posture classifications for all 1,201 participants against their accelerometer data (as in Figure 2.2 and accelerometer data plotted on the sphere (as in the bottom row of Figure 4). Problems with estimating the upright orientation or detecting change points were noted and summarized.

Second, we assess the performance of the algorithm within the Towson Accelerometer Study. In an exercise science lab, forty-four participants wore an ActiGraph GT3X triaxial accelerometer at the waist mounted with an elastic belt for one hour and were asked to perform a sequence of five minute activities: walking on a treadmill, walking on a level surface, walking while pushing a walker, fast-walking on a level surface, relaxed sitting, sitting at a table while solving a crossword puzzle, sitting while turning a hand-crank, and lying down. A research assistant recorded the start and stop times of each activity with a stopwatch. These observer-recorded labels are used as the gold standard labels.

We applied the algorithm to the resulting triaxial accelerometer data. The original data, sampled at a rate of 80 hz, was decimated to 1 hz to better reflect the sampling rate of the Zio accelerometer (1.5 hz). All the algorithm’s parameters were kept at their default values (*p*\* = 0.95, *k* = 0.98, *θ*\* = π/4, bandwidth = 0.14), but accelerometer data was classified in thirty second intervals. In the lab, it would be unreasonable to expect any participant to remove and re-position their accelerometer. Hence, to assess the accuracy of the algorithm in the case where devices were removed and replaced, an additional simulation analysis was performed. Each participant was assigned a random, uniformly distributed time that indicated when the device would be rotated. Additionally, a random rotation matrix was assigned to each participant to indicate how their data would be rotated after the simulated rotation time. Accuracy, sensitivity, and specificity were summarized and compared to Actigraph’s algorithm, described in Section 3.0. Actigraph assumes the upright orientation *μ_i_* = (0, —1, 0) and classifies a person as recumbent recumbent if the inclination is greater than 65 degrees (*θ_i_*(*t*) > 1.13).

## 4. Results

### 4.1 External Validation via the Towson Accelerometer Study

The recumbent rest algorithm was applied to the Towson Accelerometer Study triaxial accelerometer data. The experiment resulted in an average of 44 minutes of labeled activity data per participant, 5.5 minutes (12.8%) of which was spent lying down. When no simulated rotations were added to the data, our algorithm correctly labelled 96.0% of 30 second intervals as either recumbent or upright (sensitivity = 97.5%, specificity = 95.9%), and achieved 100% accuracy for 29 individuals (66%). The most common false positive was labelling relaxed sitting as lying down: 23% of relaxed sitting intervals and 5% of reading while sitting intervals were labelled recumbent. When simulated device rotations were added to the data, the algorithm had an accuracy of 93.5%, sensitivity of 97.7%, and specificity of 93.5%. In comparison to our algorithm, when no simulated rotations were added to the data, our implementation of Actigraph’s algorithm for hip-worn data gave an accuracy of 96.4% (sensitivity = 90.7%, specificity = 97.6%).

### 4.2 MACS: Zio Wear Time and Device Removal

Accelerometer files were obtained for 1, 250 participants in the MACS Zio substudy, which contained a total of 12,632 days of recording, or 10.1 days per person (median = 13.0, Q1 = 6.0, Q3 = 14.0). After screening accelerometer data for periods of non-wear or device malfunction (see Appendix 3 for details on non-wear screening), we detected 84.8 days of non-wear among 74 participants. Forty-nine participants (3.9%) had fewer than 24 hours of valid, consecutive recording after screening for non-wear. We removed non-wear from the timeseries and excluded participants who had fewer than 24 hours of consecutive wear time. The remaining 1,201 participants had 10.4 days of valid wear per person (Median = 14.0; Q1 = 6.7; Q3 = 14.0). For participants whose valid wear was interspersed by periods of non-wear, the posture algorithm was applied separately to contiguous periods of valid wear.

### 4.3 MACS: Upright Orientation of Devices

The population central upright orientation during wear periods immediately following device placement was 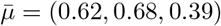 with mean resultant length 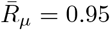. The median angular difference between this population upright orientation and each individual’s estimated initial upright orientations was 10.3 degrees (Q1 = 6.5, Q3 = 15.4). These results indicate, while there was a strong tendency for most individuals to have an upright orientation near the population center, using a common upright orientation for all individuals would be inappropriate: a 15 degree error in estimation of the upright orientation would likely induce errors in posture classifications. Using the methods described in section 3.2, we estimate that 43 participants (3.6%) removed and re-positioned their devices. We examine the sensitivity of this number to various tuning parameters in Appendix 2a and 2b.

### 4.4 MACS: Daily Time Spent Recumbent

Overall, the median daily time spent recumbent among MACS participants was 9.97 hours (Q1 = 8.53, Q3 = 11.83). Overall, 87.1% of study participants were recumbent at 4 am, and 15.5% were recumbent at 1pm. Participants were recumbent more on weekends (median = 10.4 hours/day, Q1 = 8.32, Q3 = 13.3) than on weekdays (median = 9.5 hours/day, Q1 = 7.6, Q3 = 12.1). The left panel of Figure 5 displays the estimated proportion of study participants who were recumbent at any given time of the day, and the right panel displays the proportion of the day participants were recumbent on any given day of the week.

**Fig. 5.**
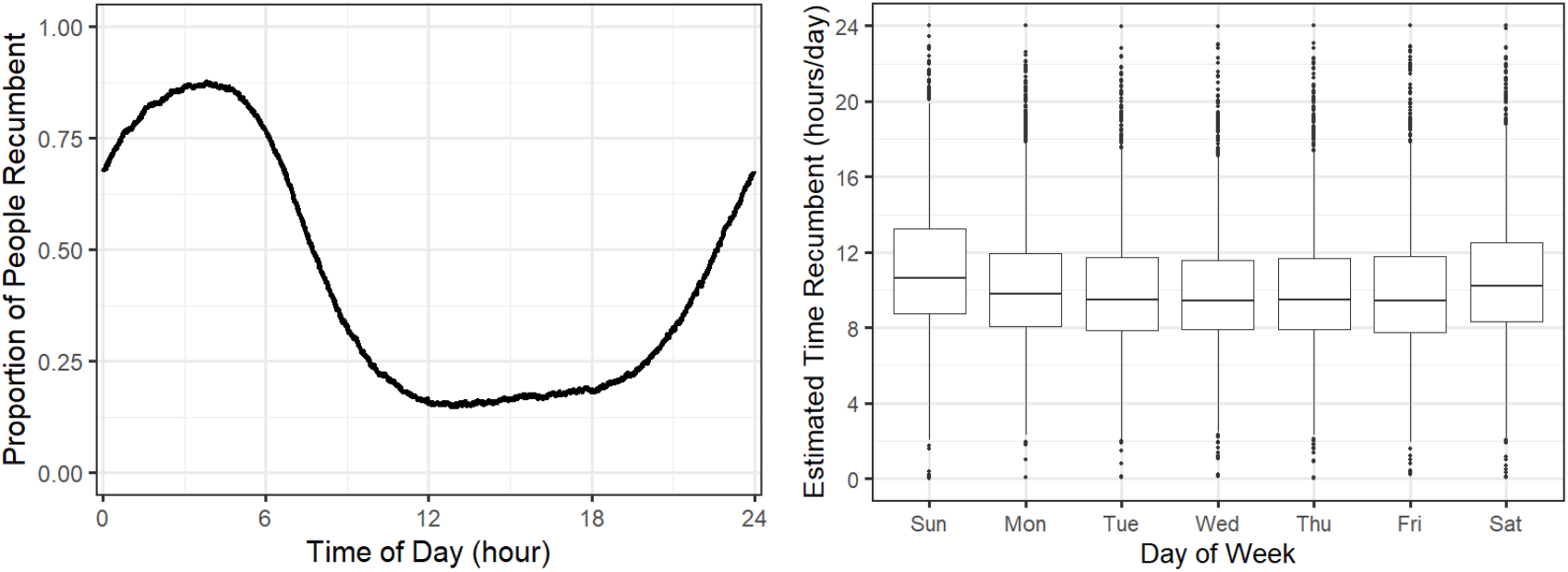
Population summaries of algorithm for classifying posture. The left panel displays the estimated proportion of participants who are by time of day. The right panel displays the distribution of the number of hours per day spent recumbent by day of the week.

### 4.5 MACS: Quality Control

During manual inspection of posture classifications for each individual, 12 individuals were found to have an improperly-estimated upright orientation (1.0%). We failed to identify the existence or timing of 9 device removal and replacements (0.7%). Three individuals likely removed and replaced their device more than once (0.25%). Hence, overall 1.95% of 1,201 participants had substantial classification errors.

## 5. Discussion

We extend existing methods for posture classification in the setting of ECG patches in large cohort studies. Our extension estimates the individual-specific upright orientation of the device without need for calibration measurements in the lab, accounts for changes in this upright orientation when participants remove and reposition their device, and utilizes clustering to generate individual-specific decision boundaries for posture classifications. En route to producing this algorithm, we discovered and demonstrated that device removal and replacement can raise important problems both for processing accelerometer and ECG data. Failing to account for device removal and replacement has the potential not only to introduce errors into posture classification algorithms but also into algorithms for processing ECG data. We confirmed that after participants had flipped their devices upside-down, the ECG signal flipped as well.

To the best of our knowledge, we also present the first published analysis of Zio accelerometer data and its potential for quantifying and studying multi-day continuous trajectories of posture. The lack of previous publications on Zio accelerometer data is likely due to lack of dedicated software from the manufacturer. The extracted accelerometer data can be studied by itself as a source of information about physical activity and behavior as well as jointly with ECG measurements. While other accelerometers have been widely used to study physical activity, we have found that utilizing Zio could be preferable in the case where a study has already collected Zio data, so obtaining Zio accelerometry would pose no new costs to site staff or participants. Additionally, a study might aim to understand ECG responses to physical activity, which is difficult to study with separate ECG and accelerometer devices due to device drift. Zio has a single internal clock which synchronizes the accelerometer and ECG monitor, making joint analysis of the data simpler.

While we have not included an example of how our posture algorithm could be used to analyze ECG data, we have previously published work that demonstrated posture was an important correlate of the QT Variability Index (QTVI) in the Multicenter AIDS Cohort Study. On average, QTVI was 0.32 lower during recumbent periods than upright periods among MACS participants (95% CI = 0.31, 0.33). This finding was similar in character to lab-based studies of QTVI and posture (Yeragani *and others*, 2000b). Our findings demonstrated the importance of adjusting for posture in analysis of ECG data, and that group comparisons can easily be confounded by differences in posture.

The most important limitation of our approach is the lack of labelled data to test our algorithm against. The algorithm was built using a set of 14 individuals, and four tuning parameters were selected according to sensitivity analyses and careful inspection of plots. Applying the algorithm in the Towson Accelerometer Study suggested it performed with accuracy, sensitivity, and specificity comparable to the algorithm used by ActiGraph. Yet, some might take issue with this approach as the Towson Accelerometer Study used a waist-mounted device rather than a chest-mounted device, so error rates from this experiment may not translate directly to the free-living data collected using Zio. We believe our algorithm would be robust to this placement. This is because the only assumptions we make about the placement of the device are (1) the device is on a body location that will tend to have a single orientation when active, and (2) when a person is recumbent the orientation of the device will be roughly perpendicular to the active orientation. While these assumptions would likely be inappropriate for devices mounted to the arms or legs, we assert that they are likely true for any location on the human trunk. Furthermore, we did not train this algorithm using data from the Towson Accelerometer Study. Rather we used it as a test: reasonable performance in this different setting would likely indicate reasonable performance for the original setting.

In conclusion, accelerometers in ECG patches pose powerful new opportunities for joint analysis of human activity and heart health. We have taken a first step toward giving researchers the tools to process accelerometer data from the Zio XT into meaningful measurements of human activity. While we have focused primarily on the classification of posture, additional research could be conducted on measurements of activity intensity and their utility in models of ECG features, adverse cardiac events, and arrhythmias.

## 6. Supplemental Material

### 6.1 Software and Vignettes

We produce an R package “postuR” that streamlines the data processing steps needed for posture classification. The goal for this package is to provide a data processing pipeline that inputs the raw accelerometer data and outputs summaries of activity and posture. The package contains the sample of raw Zio accelerometer data for one individual (Person 1 from Figure 3). The vignette for this package contains code to replicate the left column of Figure 3 from the paper.

### 6.2 Data

Zio accelerometer data was collected and is owned by the MACS study. MACS data can be applied for at https://aidscohortstudy.org/researchers/. We obtained permission to release a full accelerometry data set (with changed dates) for an individual whose data was featured in a number of visualizations in the paper. This data has been included in the R package.

Data from the Towson accelerometer study is owned and stored by Dr. Nicholas Knuth of Towson University. His contact information can be found here: https://www.towson.edu/chp/departments/kinesiology/facultystaff/nknuth.html.

### 6.3 Other Code

Data for reproducing statistics, tables, and figures from the paper and appendices will be available on Github at the time of publication.

## Acknowledgments

Work for this manuscript was supported by grants 5T32AG000247-25 from the National Institute on Aging; 5R01HL125053-04 from the National Heart, Lung, and Blood Institute; and 5R01NS060910-12 from the National Institute of Neurological Disorders and Stroke.

The authors acknowledge the valuable contributions of the other MACS investigators, staff and participants. Data in this manuscript were collected by the Multicenter AIDS Cohort Study (MACS). MACS (Principal Investigators): Johns Hopkins University Bloomberg School of Public Health (Joseph Margolick, Todd Brown), U01-AI35042; Northwestern University (Steven Wolinsky), U01-AI35039; University of California, Los Angeles (Roger Detels, Otoniel Martinez-Maza), U01-AI35040; University of Pittsburgh (Charles Rinaldo, Jeremy Martinson), U01-AI35041; the Center for Analysis and Management of MACS, Johns Hopkins University Bloomberg School of Public Health (Lisa Jacobson, Gypsyamber D’Souza), UM1-AI35043. The MACS is funded primarily by the National Institute of Allergy and Infectious Diseases (NIAID), with additional co-funding from the National Cancer Institute (NCI), the National Institute on Drug Abuse (NIDA), and the National Institute of Mental Health (NIMH). Targeted supplemental funding for specific projects was also provided by the National Heart, Lung, and Blood Institute (NHLBI), and the National Institute on Deafness and Communication Disorders (NIDCD). MACS data collection is also supported by UL1-TR001079 (JHU ICTR) from the National Center for Advancing Translational Sciences (NCATS) a component of the National Institutes of Health (NIH), and NIH Roadmap for Medical Research. Dr. Ashikaga receives research funding from The Foundation Leducq Transatlantic Network of Excellence (16CVD02). The contents of this publication are solely the responsibility of the authors and do not represent the official views of the National Institutes of Health (NIH), Johns Hopkins ICTR, or NCATS. The MACS website is located at http://aidscohortstudy.org/.

The authors acknowledge the valuable contribution of Dr. Jennifer Schrack (Johns Hopkins University School of Public Health) and Dr. Nicholas Knuth (Department of Kinesiology, Towson University) for the collection of labeled accelerometry data for validation purposes.

