## Appendix for "Classification of Free-Living Body Posture with ECG Patch Accelerometers: Application to the Multicenter AIDS Cohort Study"

**Appendix 1:** Choice of Clustering Algorithm, Tuning Parameters for Clustering

Five types of clustering were considered: hierarchical clustering (HC) methods single, ward, centroid, and median (SHC, WHC, CHC, MHC) and mean shift clustering (MSC). Initially Von Mises mixture modelling (VMMM) and density-based spatial clustering (DBSCAN) were also considered, but were quickly eliminated as they were considered inappropriate for our data: VMMM forces clusters to be circular, which seems inappropriate as demonstrated in Figure 4. While DBSCAN nicely captured some of the irregular shapes seen in our data, it does not assign every point to a cluster, instead labelling as "noise" some points at the margins of clusters.

For 14 individuals in the training data set, we first circled by hand the cluster we felt represented an upright position. We aimed to identify a clustering method and associated tuning parameter value that for the most individuals (1) separated our selection of upright points from the remaining points and (2) did not further sub-divide the selection of upright points. For HC methods, we tuned the number of clusters directly, while for MSC we varied the bandwidth over a grid of values. (Though the analyst does not determine the number of clusters directly when using MSC, a lower bandwidth tends to yield a larger number of clusters.)

Figures 1 through 5 depict the results of varying the tuning parameters for the five clustering methods for just one individual. They respectively demonstrate that SHC, CHC, MHC, and WHC achieved goal (1) after identifying 10, 5, 6, and 5 clusters, and failed goal (2) after 15, 10, 7, and 7 clusters. MSC achieved goal (1) for this individual with a bandwidth of 0.2, but did not fail goal (2) with a bandwidth as low as 0.06.

Using all individuals, the following optimal tuning parameters were chosen for each clustering method. The optimal bandwidth for MSC was 0.14 and achieved both goals (1) and (2) for 79% of participants. The optimal cluster count for CHC was 7 and achieved both goals (1) and (2) for 89% of participants. The optimal cluster count for WHC was 5 and achieved both goals (1) and (2) for 74% of participants. The optimal cluster count for MHC was 7 and achieved both goals (1) and (2) for only 26% of participants. Using up to 15 clusters, SHC failed goal (1) for 74% of participants so no parameter selection was attempted. While CHC appeared to perform better at its optimum than MSC or WHC (89% vs. 79% and 74%), we considered all three to be reasonable enough to warrant a sensitivity analysis, which is detailed in Appendix 2c.

### **Appendix 2:** Sensitivity Analyses

Our algorithm includes four tuning parameters: $p$, $k$, $\theta^*$, and the cluster count or bandwidth parameter for the chosen clustering algorithm. Introduced in Equation 3.1, $p$ determines the which time points are considered to have high accelerations, and consequently $p$ regulates how a participant's upright orientation is estimated. In section 3.2, we use $k$ to decide whether a participant's data exhibits device removal and replacement. A lower value for $k$ indicates more people will be classified as likely having removal and replacement. In Equation 3.6, we introduce $\theta^*$, the angular

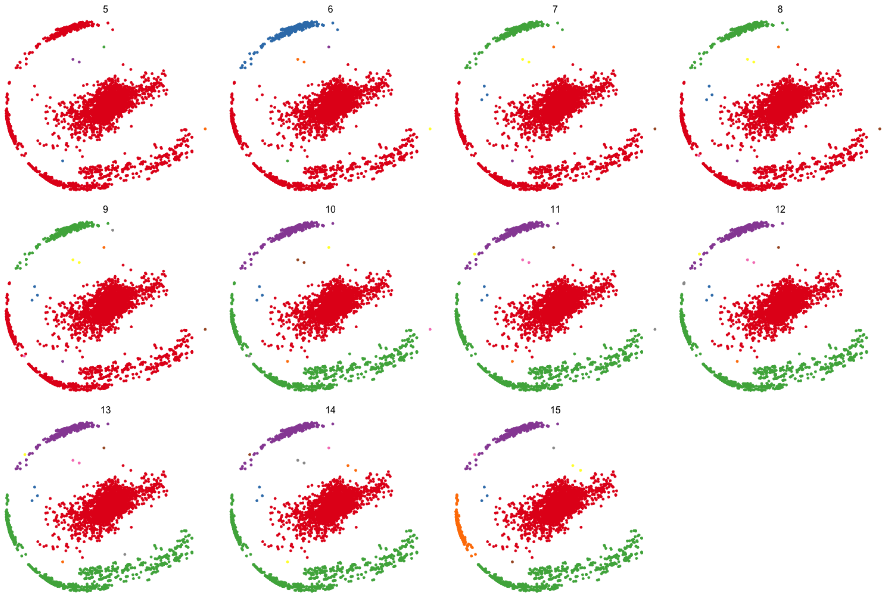


Fig. 1. Single Hierarchical Clustering tree for one individual, cut at various cluster counts (5...15). Within each plot, points within a cluster have the same color, though colors may not be comparable across plots. Here, the cluster at the center of the data (red) seems separated from clusters around the perimeter after cutting the tree at 10 or more clusters. The upright orientation does not become substantially subdivided with 15 or fewer clusters.

threshold at which we label a particular time interval recumbent or upright based on its estimated inclination. Increasing $\theta^*$ will directly reduce the number of points labelled recumbent. The last tuning parameter is the clustering algorithm and its respective tuning parameter we use to time points into posture classes. In the previous section, we showed that Mean Shift Clustering, Centroid Hierarchical Clustering, and Ward Hierarchical Clustering all worked reasonably well on a set of 14 participants. In the next four subsections, we detail the sensitivity analyses we used to choose default values for these parameters.

#### *(2a) Sensitivity to Percentile for Estimation of Upright Position*

We examined how the estimated upright orientation $\tilde{\mu}_i$ changed when we varied the percentile ($p = 0.95$) used to define the participants' individual quantiles $q_i$ of high acceleration points. We varied $p$ over the grid $0.9, 0.91 \ldots 0.99$ and for each study participant calculated (1) the pairwise

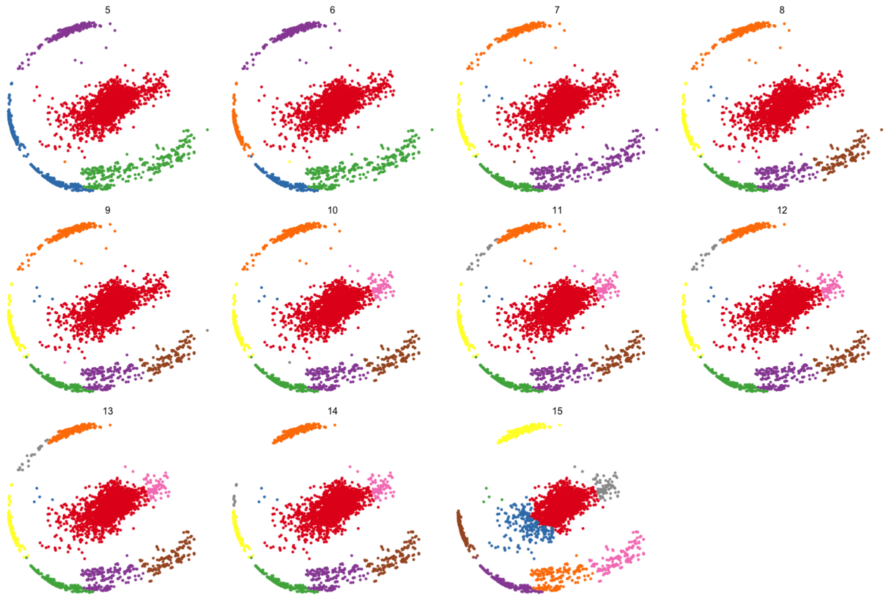


Fig. 2. Centroid Hierarchical Clustering tree for one individual, cut at various cluster counts (5 ... 15). Within each plot, points within a cluster have the same color, though colors may not be comparable across plots. Here, the cluster at the center of the data seems separated from clusters around the perimeter when cutting the tree at 5 or more clusters. The center cluster does not become substantially subdivided until 10 clusters are selected.

angular difference in the estimated upright orientation for and (2) the pairwise concordance of posture classifications. Both (1) and (2) are plotted below using $p = 0.90$ for the reference upright orientation and posture classifications. Each set of connected points represent one of thirteen of sixteen individuals who had no suspected device removal.

The left panel of Figure 6 illustrates that changing the percentile from 90% to 95% changed estimated upright orientations changed by less than $3.75^o$ for all individuals. Increasing the percentile from 90% to 99% had a changed the estimated upright orientations less than $11^o$ for all individuals.

The right panel of Figure 6 demonstrates that increasing the percentile from 90% to 95% changed less than 0.75% of the classifications per study participant (Med = 0.09%, IQR = [0.06%, 0.36%]). Furthermore, increasing the percentile from 90% to 99% changed up to 2.6% of the classifications per participant (Med = 0.55%, IQR = [0.29%, 0.92%]).

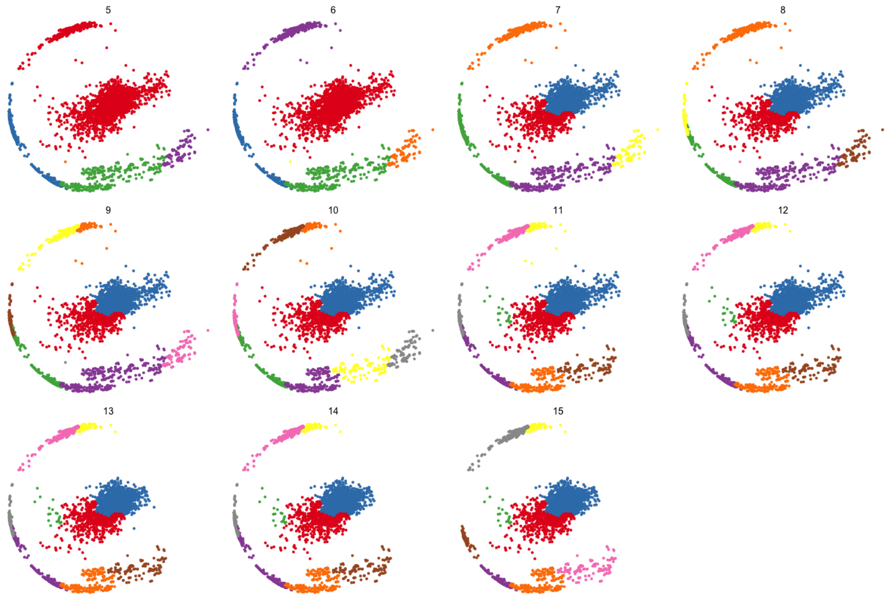


Fig. 3. Median Hierarchical Clustering tree for one individual, cut at various cluster counts (5 ... 15). Within each plot, points within a cluster have the same color, though colors may not be comparable across plots. Here, the cluster at the center of the data seems separated from clusters around the perimeter when cutting the tree at 6 or more clusters. The center cluster does not become subdivided until 7 clusters are selected.

Among MACS participants who had no evidence of non-wear or device removal and replacement (n = 1,109), increasing the threshold from $p = 0.95$ to $p = 0.99$ changed individuals' estimated upright orientations by a median of $2.18^o$ (IQR $= 1.29^o, 3.58^o$) and decreasing the threshold to $p = 0.9$ changed individuals' estimated upright orientations by a median of $2.40^o$ (IQR $= 1.20^o, 6.25^o$).

We believe this sensitivity analysis affirms our use of $p = 0.95$ to define an individuals high acceleration points, as values in a fairly wide neighborhood around 0.95 result in similar classifications. Hence, there would be only marginal benefit to optimizing our algorithm against a training data set.

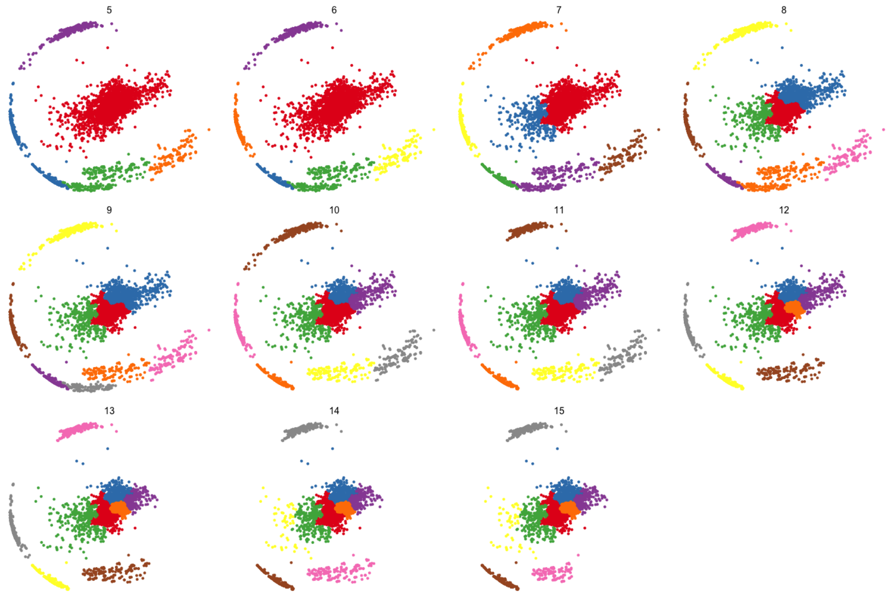


Fig. 4. Ward Hierarchical Clustering tree for one individual, cut at various cluster counts (5 ... 15). Within each plot, points within a cluster have the same color, though colors may not be comparable across plots. Here, the cluster at the center of the data seems separated from clusters around the perimeter when cutting the tree at 5 or more clusters. The center cluster does not become subdivided until 7 clusters are selected.

#### *(2b) Detecting Changes in Upright Orientations: Sensitivity to $k$ and $p$*

We examine how the people estimated to have removed and replaced a device varied with $p$ (the percentile we use to define high-acceleration points) and $k$ (the threshold for $r_i^*$ we use to label whether an individual removed and replaced their device). The plot below depicts $r_i^*$ across values of $p$.

Each point on the plot represents ratio of the within vs. marginal mean resultant length ($r_i^*$) for all individuals and all values of $p = 0.9 \ldots 0.99$. Ratios for a given individual are connected with lines. The cutoff $k = 0.98$ for labelling whether an individual removed and replaced their device is represented as a horizontal dashed line.

For a cutoff of $k = 0.98$, we see that whether we label an individual as having removed and replaced their device is independent of $p$: none of the lines cross the dashed line that represents

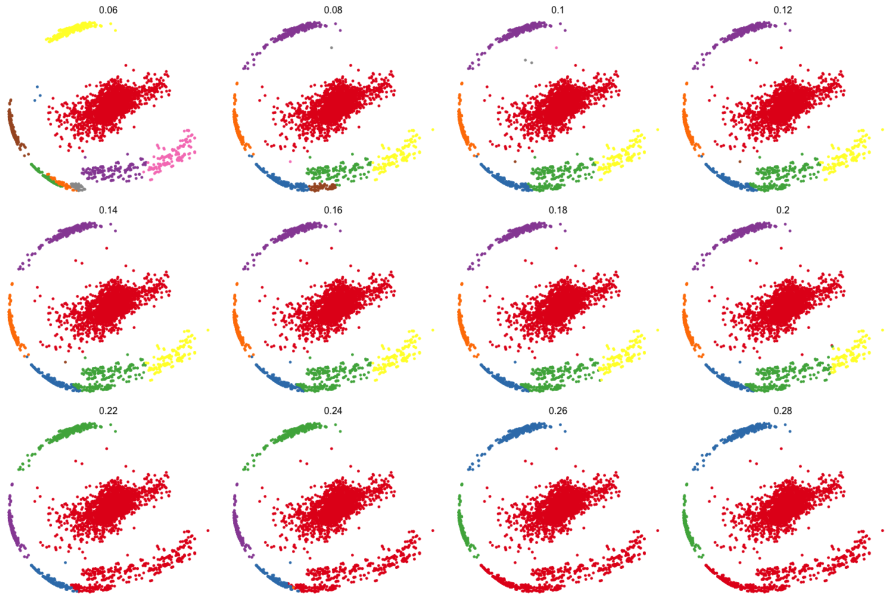


Fig. 5. Mean Shift clustering for One individual, using various bandwidths (0.06 ... 0.28). Here, the cluster at the center of the data seems separated from clusters around the perimeter when using a bandwidth 0.2 or less, resulting in 6 or clusters. The center cluster isn't divided by reducing the bandwidth to 0.06.

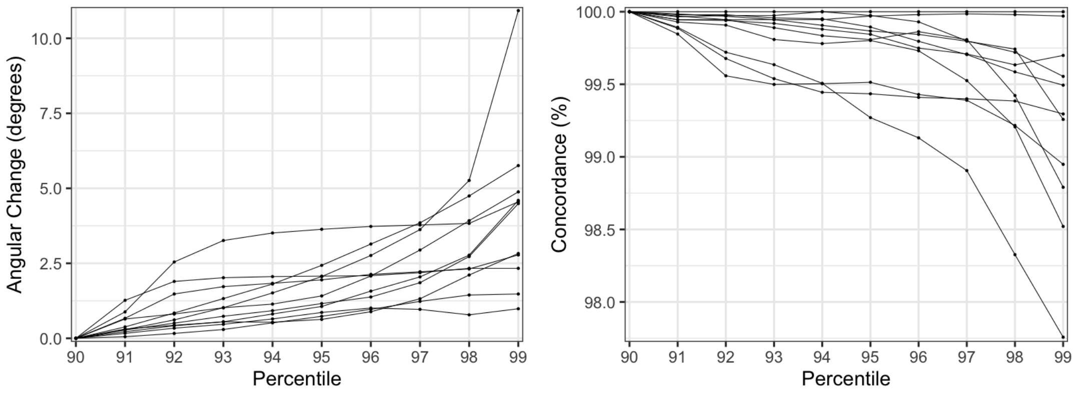


Fig. 6. Sensitivity of estimated upright orientation and posture classification to change in $p$, the percentile used to define high acceleration points.

the cutoff $k = 0.98$. We also see, regardless of $p$ none of the labels would not change until $k$ were decreased to 0.94.

This plot demonstrates that (1) a cutoff of $k = 0.98$ separates the individuals with suspected device removal/replacement regardless of the value $p$ we choose and (2) that values of $p$ in the

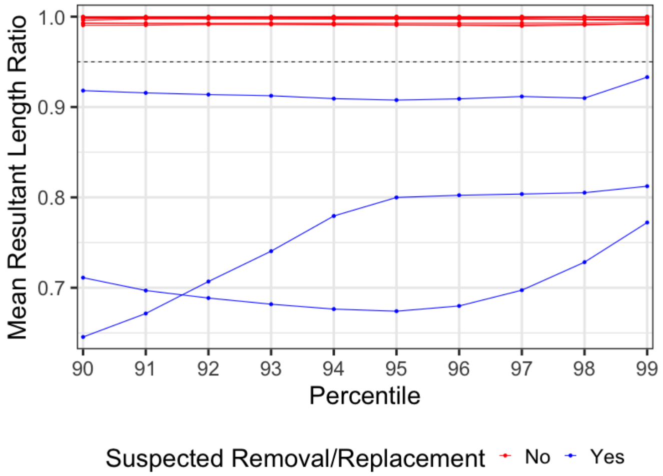


Fig. 7. Sensitivity of ratio of within vs. marginal mean resultant length ($r_i^*$)to changes in $p$

neighborhood of $p = 0.99$ might give us decreased ability to label individuals who removed and replaced their devices (note that the three ratio trajectories of individuals with suspected removal/replacement trend upwards toward the cutoff when $p = 0.99$).

among all MACS participants who had no flagged non-wear (n = 1,178). Figure XXX depicts how PRR varies with $p$ and $k$.

By decreasing $p$ from 0.95 to 0.9, we include lower-acceleration movement in the estimation of the upright position, and Figure XXX shows PRR is increased for all values of $k$.

However, Figure XXX also demonstrates that increasing $p$ from 0.95 to 0.99 only substantially decreases PRR when $k \geqslant 0.96$).

#### *(2c) Sensitivity to Choice of Clustering Algorithm*

As stated in Appendix 1, Mean-Shift Clustering (bandwidth = 0.14), Centroid Hierarchical Clustering (7 clusters), and Ward Hierarchical Clustering (5 clusters) performed reasonably among the individuals on which the algorithm was built. We tabulate the individual-specific concordance of classifications between these three methods below.

Table 1. Individual-Specific Concordance of Classifications between Three Clustering Methods

| | **Concordance** | | |
|---|---|---|---|
| **Method** | **Median** | **Q1** | **Q3** |
| Mean Shift vs. Centroid | 99.4% | 98.2% | 99.7% |
| Mean Shift vs. Ward | 99.8% | 99.5% | 100.0% |
| Centroid vs. Ward | 98.5% | 99.3% | 99.9% |

Table 1 indicates that Concordance between the classifications between all three methods were high among the individuals in the training data set. We believe these results indicate that these three methods produce similar classifications, and the choice of clustering among these three methods should be driven by which is fastest.

#### ***(2d)*** *Sensitivity to Threshold of* $45\deg$ *for Labelling Points Recumbent*

Since an inclination of 0 degrees represents a perfectly upright posture, and 90 degrees represents a perfectly recumbent posture, a 45 degree threshold separates the points which are closer to an upright or recumbent posture. Hence, we chose a 45 degree threshold to label each point as recumbent or upright. However, we the sensitivity of our classifications to the choice of this threshold within the range 30 to 60 degrees. The results are depicted below in Figure 8.

The y axis in each panel of Figure 8 represents the overall percentage of time an individual spent recumbent, while the x axis represents the chosen angular threshold. The left panel represents the initial classifications before cluster-based majority voting, while the right panel represents the final classifications after the cluster-based majority voting. The blue lines and ribbons represent medians and IQRs. Expectedly, the estimated proportion of time recumbent decreases as the angular threshold increases. The decrease is much more gradual in the right panel, where a point's classification depends only on it's inclination, than in the left panel, where a point's classification depends on the median inclination of it's entire cluster.

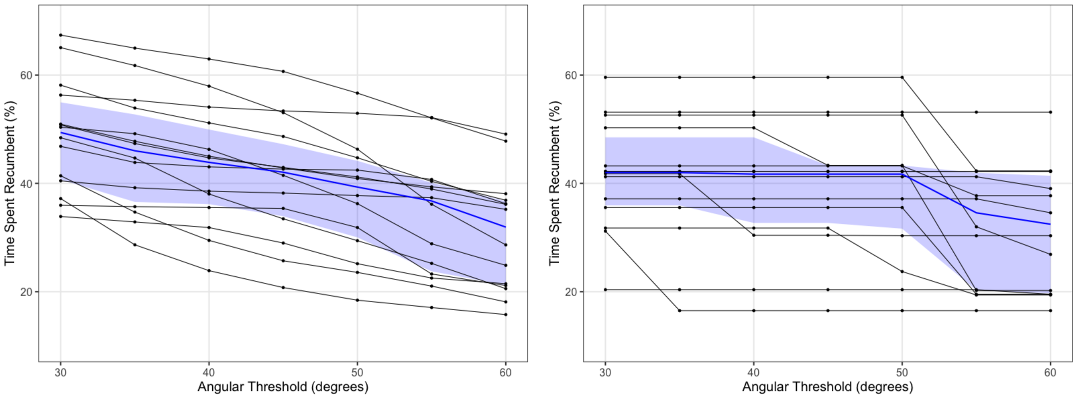


Fig. 8. Sensitivity of daily time spent recumbent to changes in angular threshold for initially classifying a point recumbent.

This plot demonstrates that–when using the cluster-based majority voting–the classifications do not change much when decreasing the angular threshold toward 30 degrees. At a threshold of 45 degrees the median percentage time spent recumbent was 42.0% (IQR $= 32.4\%, 47.3\%$), but decreasing the angular threshold to 30 increased the median percentage time spent recumbent to 43.7% (IQR $= 35.5\%, 49.3\%$). This small change indicates that the choice of an angular threshold in the interval $[30, 45]$ is likely arbitrary.

In contrast, increasing the angular threshold to 60 degrees decreased the median percentage time recumbent more dramatically to 31.6% (IQR $= 20.6\%, 37.5\%$). We believe that such a high angular threshold would be inappropriate. Figure 4 from the main manuscript demonstrated that, when looking at the postures on the sphere, some posture clusters that were seemingly recumbent could approach the 45 degree threshold from below.

### **Appendix 3:** DETECTING DEVICE NON-WEAR

A-priori, we were not sure whether non-wear periods would be evident in the data, as Zio uses an adhesive to attach to study participants and may not re-attach well to the skin. While no non-wear periods were evident among the subjects in our training data set, there were non-wear periods in the remaining MACS sample. Since we consider non-wear detection to be a device-specific procedure, we have included our methods for detecting non wear in this appendix rather than the main manuscript.

To detect non-wear in our data, we adapt the methods used by the National Cancer Institute for sub-second accelerometer data. We scan the data for three hour windows in which few changes are observed along each of the x, y, and z axes. If a point in time is contained in a three hour window in which fewer than 10 measurements represent changes, we consider that point to be in a non-wear period.

Mathematically, let $x_{it}$, $y_{it}$, and $z_{it}$ denote the triaxial timeseries observed for person $i$ at the time points $t = 1 \dots T_i$, which are equally-spaced at approximately 1.56 hz. We define a change along any axis at time $t$ as $\delta_{it} = 1 - I\{x_{it} = x_{(it+1)}\} \cdot I\{y_{it} = y_{(it+1)}\} \cdot I\{z_{it} = z_{(it+1)}\}$. Then we define $n_{it}$ as the number of changes that occurred in the three hours following time t:

$$n\delta_{it} = \sum_{(s=t)}^{s+\lfloor 180 \cdot 60 \cdot 1.561 \rfloor} \delta_{is}$$

Finally, we define $nw_{it}$, our non-wear indicator for time $t$, as

$$nw_{it} = \prod_{s=t-\lfloor 180 \cdot 60 \cdot 1.561 \rfloor}^{t} I\{nd_{it} \leqslant 10\}.$$

For all participants who our algorithm flagged as having non-wear, raw triaxial accelerometer data was plotted and manually inspected to verify that a non-wear period occurred.
